# Enforced E-selectin ligand installation enhances homing and efficacy of adoptively transferred T cells

**DOI:** 10.1101/2025.01.12.632650

**Authors:** Yingqin Hou, Jinfeng Ye, Ke Qin, Leigh-Ann Cui, Shereen Chung, Digantkumar Gopaldas Chapla, Wenjian Wang, Yujie Shi, Mingkuan Chen, Kelley. W. Moremen, Robert Sackstein, Peng Wu

## Abstract

Adoptive T-cell transfer has revolutionized the treatment of hematological malignancies. However, this approach has had very limited success in treating solid tumors, largely due to inadequate infiltration of vascularly administered T cells at tumor sites. The shear-resistant interaction between endothelial E-selectin and its cognate ligand expressed on leukocytes, sialyl Lewis X (sLe^X^), is an essential prerequisite for extravasation of circulating leukocytes. Here, we report that enforced E-selectin ligand expression (enforced sLe^X^ display) on antigen-specific T cells can be achieved by fucosylating cells via cell surface treatment with the human α1-3-fucosyltransferase, FUT6 (“exofucosylation”), or via Golgi-targeted FUT6 overexpression (“Golgi-fucosylation”). However, despite comparable E-selectin binding, only sLe^X^-modified T cells engendered by exofucosylation, not by Golgi-fucosylation, exhibited enhanced parenchymal infiltration of target malignant sites. This heightened homing yielded significantly improved therapeutic efficacy in various murine syngeneic and xenograft cancer models, including subcutaneous solid tumors, lymphoma and leukemia, as well as lung and bone marrow metastases. Therefore, exofucosylation represents a promising strategy to improve the efficacy of adoptive T-cell therapy, particularly in the treatment of solid tumors and metastatic disease.

## Introduction

An unresolved challenge in adoptive cell therapy (ACT) is to channel adoptively transferred immune cells specifically to their target sites and thereby avoid non-specific tissue migration. After intravenous administration, the majority of transferred T cells are sequestered in non-tumorigenic tissues, initially accumulating in the lungs, followed by the liver and spleen.^1,2^ Only a small fraction of these cells ultimately reaches the intended tumor sites, which not only limits therapeutic efficacy, but also poses safety risks. As a result, extensive *in vitro* expansion is often required to generate clinically relevant quantities of T cells for infusion into patients. This process is associated with high cost^3^ and can also lead to T cell exhaustion, further compromising their persistence and functional efficacy *in vivo*. Even when T cell exhaustion is avoided, infusing large numbers of activated T cells increases the likelihood of severe adverse effects, such as cytokine release syndrome (CRS).^4,5^ These limitations underscore the need for strategies to enhance T cell target-specific trafficking to maximize the therapeutic potential of adoptive cell therapies.

Efficient homing, whereby immune cells selectively colonize target tissues, is critical to the success of ACT. Homing begins with the tethering and rolling of blood-borne cells on the target endothelium, a process mediated primarily by the interaction of E-selectin with its cognate glycan ligand, sialyl Lewis X (sLeX, CD15s)^.6,7^ E-selectin, an endothelial lectin constitutively expressed in bone marrow and skin microvessels,^8^ is upregulated in nearly all microvascular endothelial beds in response to inflammatory cytokines such as TNFα and interleukin-1ß^.9,10^ Notably, E-selectin is also expressed in the tumor microenvironment,^11^ although its expression patterns in different solid tumors remain underexplored.

sLe^X^ is a tetrasaccharide composed of sialic acid and fucose attached to the most common glycan epitope, *N*-acetyllactosamine (LacNAc), at the periphery of the leukocyte glycocalyx^.12,13^ While the expression of this glycan epitope on genetically engineered T cells, including TCR-T and chimeric antigen receptor (CAR)-T cells, expanded *in vitro* has been poorly characterized, chemoenzymatic fucosylation can be applied to increase its expression.^14,15^ In this approach, a recombinant α1-3 fucosyltransferase is utilized to convert type 2 sialyl LacNAc to sLeX directly on the cell surface, offering a simple procedure with the potential to be seamless integrated into therapeutic T cell manufacturing pipelines. However, the impact of enforced sLeX expression on the efficacy of the modified T cells for cancer treatment has produced conflicting results. On the one hand, two studies have shown that enforced sLeX expression enhances T-cell homing to the bone marrow^,14,16^ with one of these studies also reporting improved tumor control^16^. On the other hand, a separate study found that cell surface fucosylation provided minimal benefits for CAR-T cell tissue homing and therapeutic efficacy.^17^ To date, a clear causal relationship between enforced E-selectin ligand expression and the ability of adoptively transferred T cells to infiltrate the tumor microenvironment and generate productive therapeutic efficacy has not been established.

Here, we use cell-surface direct fucosylation mediated by human α1-3 fucosyltransferase (FUT6) or its Golgi-targeted overexpression to enforce E-selectin ligand presentation on therapeutic T cells, including TCR-T and CAR-T cells. We will then evaluate the functional impact of these modifications through adoptive transfer of the engineered T cells into various murine cancer models, allowing a comprehensive assessment of enhanced E-selectin ligand display in improving T cell tumor homing and anti-tumor immunity.

## Results

### Enforced sLe^X^ expression on T cells improves tumor-specific homing

We chose recombinant human α1-3-fucosyltransferase, FUT6, as the enzyme to enforce the expression of E-selectin ligand, sLe^X^, on the cell surface of T cells due to its high enzymatic activity. To determine the optimal condition for cell-surface fucosylation, we used FUT6-mediated fucosyl-biotinylation of OT-I T cells as the model system, in which guanosine 5’-diphospho-β-L-fucose conjugated with biotin (GF-Biotin) was utilized as the donor substrate for easy detection by fluorescently labeled streptavidin. OT-I T cells express a transgenic T cell-receptor (TCR) specific for the SIINFEKL peptide (OVA_257-264_) derived from chicken ovalbumin (OVA) in the context of the MHC-I molecule H2-K^b^ (Figure 1A-B). As reported by us previously, upon antigen-stimulation and expansion with IL-2 or IL-7/IL-15, activated OT-I T cells express high levels of acceptor substrates for fucosylation.^18^ As expected, a dose-dependent addition of biotinylated fucose on T cells was achieved and the labeling signal leveled off at 200-500 uM GF-Biotin when FUT6 was used in the range of 0.05 to 1 mg/mL(Figure 1B). Therefore, we chose 0.05 mg/mL of FUT6 and 500 uM GDP-fucose for the subsequent cell-surface fucosylation studies. Under these conditions, fucosylated OT-I cells had significantly increased levels of sLe^x^ compared to control OT-I cells (Figure S1A), resulting in markedly enhanced E-selectin binding (Figure 1C). Interestingly, fucosylated OT-I T cells also grew slightly faster than control, unmodified T cells (Figure S1B, S1C), and exhibited a stronger anti-tumor activity at different effector:tumor ratios (Figure 1D and Figure S1D).

**Figure 1.**
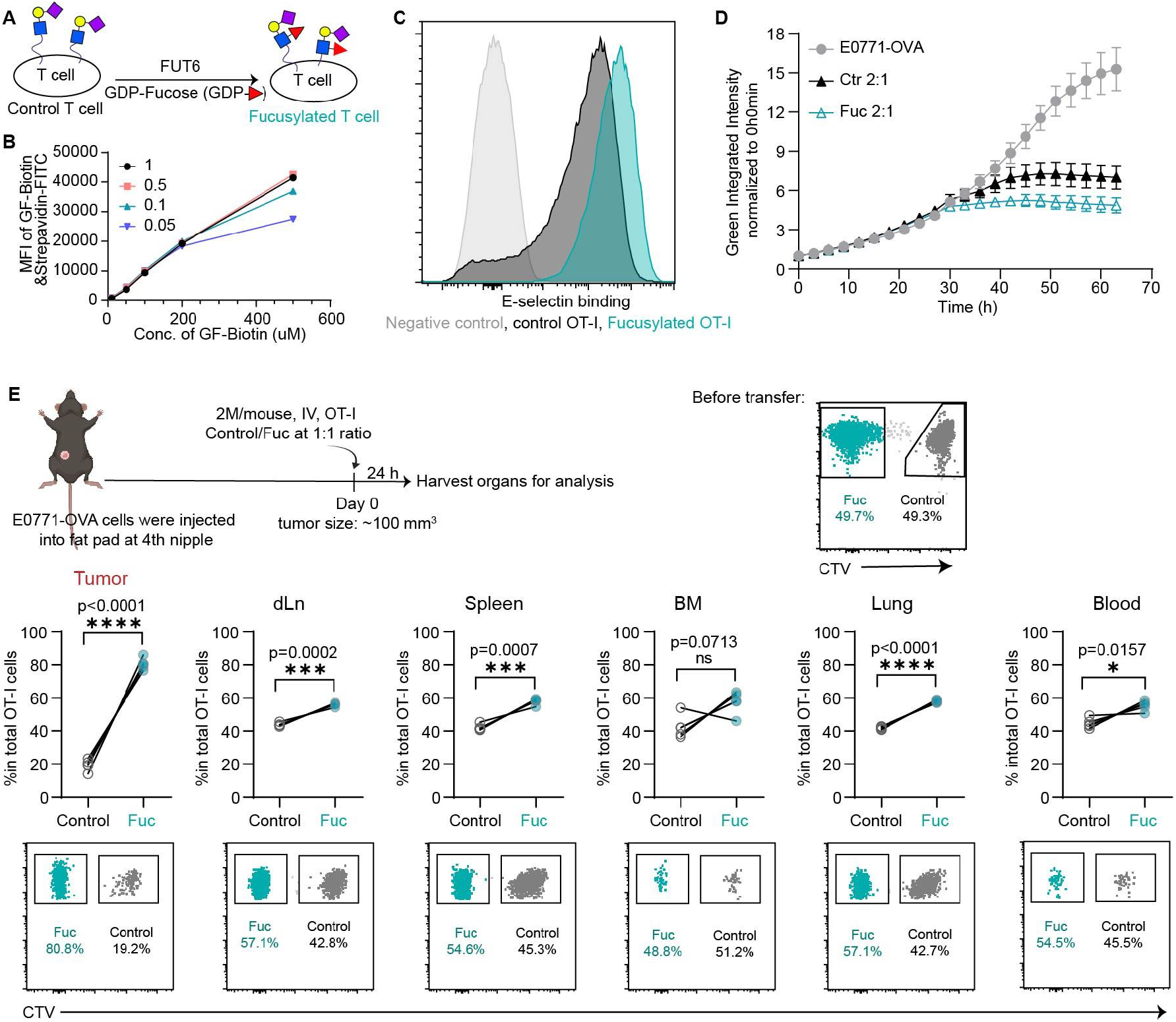
Cell-surface fucosylation on OT-I cells enhances E-selectin binding and improves specific homing to E0771-ova tumors. (**A**) Schematic workflow for cell-surface fucosylation. (**B**) Fucosylation with different concentrations of donor substrate GDP-Fucose Biotin (GF-Biotin) (0-500 uM) and fucosyltransferase as indicated in the figure (mg/mL). (**C**) E-selectin binding of control and fucosylated OT-I cells expanded with IL-2. (**D**) *In vitro* anti-tumor activity of control (Ctr) and fucosylated (Fuc) OT-I cells at an effector/tumor ratio of 2:1. (**E**) Unmodified OT-I cells labeled with Cell TraceViolet (CTV) and fucosylated OT-I cells (CTV-negative) were mixed in a 1:1 ratio and adoptively transfered into mice bearing orthotopically inoculated E0771-OVA. Organ distribution of the transferred cells was analyzed 24 hours after adoptive transfer. IV, intravenously injection. 3 repeats for *in vitro* and 5 repeats for *in vivo* studies in each group (n=3 or 5), nsP > 0.05; *P < 0.05; **P < 0.01; ***P < 0.001; ****P < 0.0001; analyzed by student T-test and mean ±standard deviation (SD) values of biological replicates.

Next, to evaluate the influence of fucosylation on T cell trafficking *in vivo*, we transferred control unmodified and fucosylated OT-I T cells into tumor-free mice separately and analyzed their organ distribution 24h after adoptive transfer. Fucosylated OT-I cells showed better homing ability to all analyzed organs, including the spleen, lung, bone marrow and inguinal lymph node, than control OT-I cells (Figure S1E). Given these findings, we next sought to determine how the presence of tumors and tumor-associated E-selectin expression might alter T cell trafficking patterns. To this end, we then examined the E-selectin expression patterns in various commonly used implantable murine solid tumor models and found that most orthotopically and subcutaneously implanted tumors, except B16 melanoma, possess high levels of E-selectin expression (Figure S1F). To directly compare the homing ability of OT-I cells with or without cell-surface fucosylation, we co-transferred control unmodified OT-I cells labeled with Cell TraceViolet (CTV) and fucosylated OT-I cells (CTV-negative) in a 50%:50% cell ratio into mice bearing orthotopically inoculated triple negative breast cancer E0771 cells engineered to express OVA (E0771-OVA). Various organs were harvested 24 hours after adoptive transfer (Figure 1E). To our surprise, only slightly higher ratios of fucosylated OT-I cells than control OT-I cells were found in most tissues examined, including the draining lymph node (dLn), spleen, lung and blood, whereas almost four times more fucosylated OT-I cells than control OT-I cells were found in the tumor tissue. Taken together, enforced expression of sLe^x^ via FUT6-mediated cell-surface fucosylation significantly enhanced the colonization of T cells in solid tumors with detectable levels of E-selectin expression.

### Cell-surface enforced E-selectin ligand expression improves CAR-T cell efficacy for treating solid tumors by enhancing T cell homing

Enhanced T cell homing to tumor tissues is expected to translate into improved tumor control. To evaluate this possibility, we adoptively transferred mouse CAR-T cells targeting human carcinoembryonic antigen (hCEA) into mice harboring subcutaneously implanted colorectal cancer cells, MC38, engineered to express hCEA (MC38hCEA). Notably, elevated E-selectin expression was readily detected in the tumor microenvironment of this model (Figure S1F), supporting its potential role in facilitating T cell homing. To investigate the impact of cell-surface fucosylation on CAR-T cell tumor homing, control CAR-T cells (labeled with CTV) were mixed with fucosylated CAR-T cells (CTV negative) in a 1:1 ratio and co-transferred into MC38hCEA-bearing mice (Figure 2A). After 24 hours, approximately equal numbers of control and fucosylated CAR-T cells were found in the blood, liver, lung and spleen. Interestingly, slightly higher numbers of fucosylated CAR-T cells were detected in the bone marrow (BM) and dLn, (∼60% vs 40-45%). The most striking difference, however, was found in the tumor where fucosylated CAR-T cells outnumbered their control counterparts by a factor of four (∼80% vs 20%). To determine whether CAR-T cells had extravasated into the tumor tissue from the blood vasculature at the time of organ harvest, tumor-bearing mice were injected intravenously with a fluorescently (phycoerythrin, PE) labeled anti-mouse CD45.2 antibody 3 minutes prior to euthanasia (Figure 2C). This approach allows differentiation between CAR-T cells circulating in the bloodstream or adhering to the vascular endothelium, which would be antibody-labeled, and those that had extravasated into the tumor tissue, which would remain labeling-free. Flow cytometry analysis revealed that only 1% of the transferred CAR-T cells were labeled, while ∼99% of CAR-T cells were protected from the labeling (Figure 2D), indicating that the vast majority of CAR-T cells had successfully infiltrated the tumor tissue. These findings were further confirmed by immunofluorescence imaging of tumor sections (Figure 2E and Figure S2). Consistent with improved tumor homing, fucosylated CAR-T cells induced superior tumor control and significantly prolonged survival in recipient mice (Figure 2F), with complete tumor eradication achieved in two of the treated mice (Figure 2G).

**Figure 2.**
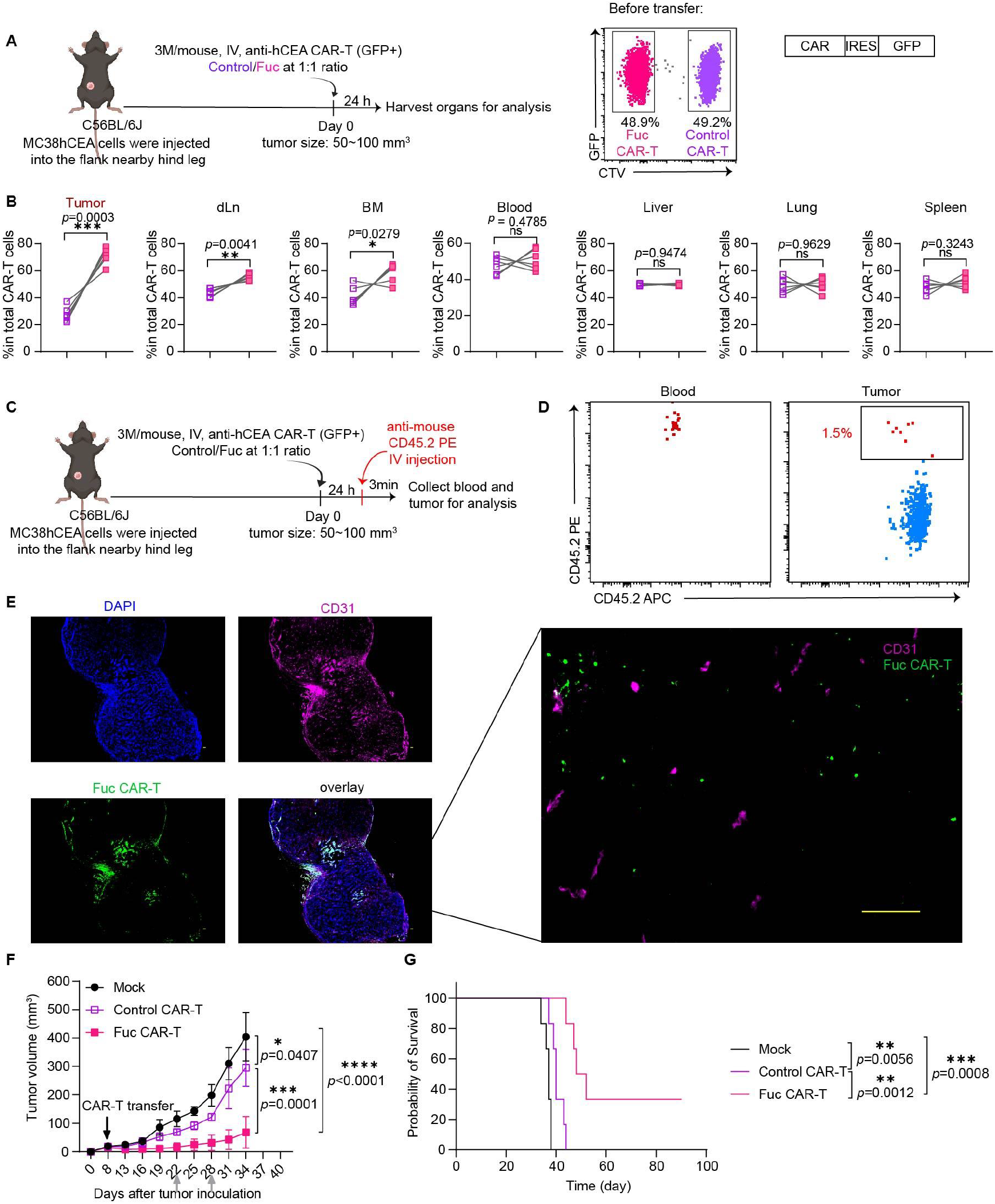
Cell-surface fucosylation improves the efficacy of CAR-T cell therapy by enhancing T cell colonization in solid tumor. **(A)** Experimental setup. (B) Tissue distribution of the adoptively transferred anti-hCEA mouse CAR-T cell at 24h after adoptive transfer (the percentages of control vs. fucosylated cells in total CAR-T cells in the tissue were shown). (C) Experimental setup and (D) flow cytometry analysis to distinguish transferred T cells that have entered tumor parenchyma from T cells in blood vasculature. (E) Distribution of fucosylated CAR-T cells in the MC38hCEA tumor analyzed by immunofluorescence, tumor #1, scale bar: 100 μm. *In vivo* antitumor efficacy (F) of CAR-T cells with or without fucosylation against MC38-hCEA colon cancer and mouse survival (G). 1×10^6^ MC38hCEA tumor cells were subcutaneously injected into the C57BL/6J mice and 0.7×10^6^ CAR-T cells were transferred into the recipient mice on day8 after tumor inoculation with lymphocyte depletion by irradiation with a dose of 5 Gy before CAR-T cell transfer. IV, intravenously injection. Grey arrows indicate the intraperitoneal anti-mouse IL-6 injection (150 ug/mouse). 5-6 repeats for each group (n=5-6), nsP > 0.05; *P < 0.05; **P < 0.01; ***P < 0.001; ****P < 0.0001; survival curves were analyzed by log-rank test and other analysis used student T-test, and mean ±standard deviation (SD) values of biological replicates.

To confirm the role of E-selectin expression in mediating enhanced tumor homing and efficacy, we utilized the B16E5 melanoma model, which expresses a chimeric murine EGFR with six amino acid mutations to allow binding of the anti-human EGFR antibody cetuximab. Unlike the MC38hCEA model, the B16E5 model has negligible E-selectin expression in the tumor microenvironment (Figure S1F). Following adoptive transfer, comparable numbers of fucosylated and unmodified anti-EGFR CAR-T cells were observed in the tumor parenchyma. Furthermore, fucosylated CAR-T cells fail to provide improved tumor control compared to unmodified CAR-T cells (Figure S3). These findings indicate that the enhanced tumor-specific homing and the resulting antitumor efficacy improvement conferred by T-cell surface fucosylation is critically dependent on the E-selectin expression in the tumor microenvironment.

### Enforced E-selectin ligand expression boosts T-cell homing to tumor metastases and strengthens anti-tumor immunity

Approximately 90% of all cancer deaths are caused by the metastatic spread of primary tumors, in which malignant cells cross physical barriers, disseminate, and colonize distant organs.^19^ In the bone vascular niche, E-selectin is known to drive mesenchymal–epithelial transition and Wnt activation in cancer cells to promote bone metastasis.^20^ Likewise, in lung metastasis, monocytes have been shown to facilitate the transendothelial migration of tumor cells by inducing E-selectin–dependent endothelial retraction and vascular permeability.^21^ Accordingly, upregulation of E-selectin has been observed in the vicinity of metastatic tumor cells in the lung.^11^ By hijacking these E-selectin-mediated processes, we hypothesized that enforced E-selectin ligand installation on T cells could enhance their migration to metastatic sites, enabling targeted elimination of disseminated tumor cells and reducing the metastatic burden.

To test this hypothesis, we used adoptive transfer of OT-I cells into the bone marrow and lung metastasis models of the triple negative breast tumor E0771-OVA. To establish experimental bone marrow metastasis, E0771-OVA cells engineered to express luciferase (E0771-OVA-luc) were injected via the caudal tail artery^22^ and the establishment of bone metastasis was confirmed by bioluminescence imaging on day 7 (Figure 3A). At 24 hours after adoptive transfer, approximately two times more fucosylated OT-I cells than control OT-I cells were found in the bone marrow. By contrast, no apparent differences were detected in other organs except for a modestly higher accumulation of fucosylated OT-I cells in the inguinal lymph nodes (Figure 3B).

**Figure 3.**
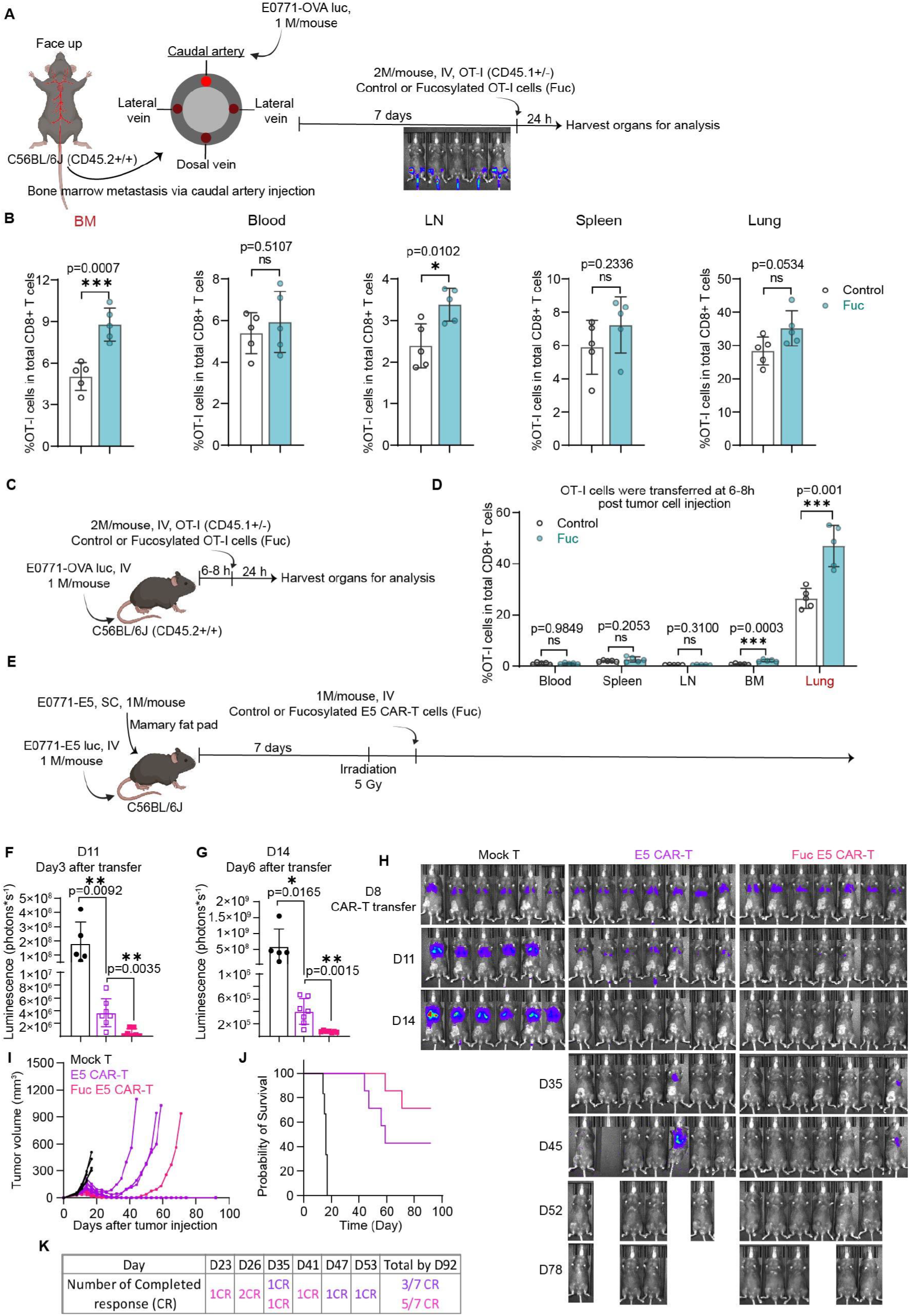
Cell-surface fucosylation enhanced CAR-T cells antitumor efficacy in both metastasis and solid tumor. (A) Bone marrow metastasis was established by tail caudal artery injection, and control or fucosylated OT-I cells were transferred into metastasis bearing mice. (B) OT-I cell number in different organs at 24h after the cell transfer. (C) Control or fucosylated OT-I cells were transferred into the mice at 6-8h after challenged by E0771-OVA cells by IV injection and (D) OT-I cell number in different organs were analyzed at 24h after the cell transfer. (E-H) Cell-surface fucosylation improves CAR-T cell efficacy by enhancing T cell infiltration in E0771-E5 tumor. 1×10^6^ E0771-E5 tumor cells were subcutaneously injected into the C57BL/6J mice and 1×10^6^ E0771-E5 expressing luciferase were intravenously injected into the same mice. CAR-T cells were transferred into the mice on day8 after tumor injection with lymphocytes deletion by irradiation with a dose of 5 Gy before the CAR-T cell transfer. IV, intravenously injection. Lung metastasis was tracked by IVIS (E-H), tumor size (I), survival curves (J) and complete response (K) were recorded. 5-7 repeats for each group (n=5-7), nsP > 0.05; *P < 0.05; **P < 0.01; ***P < 0.001; ****P < 0.0001; tumor size was analyzed by student T-test and survival curves were analyzed by log-rank test. Mean ±standard deviation (SD) values of biological replicates.

These encouraging results prompted us to further evaluate the effectiveness of fucosylated OT-I cells in the lung metastasis model. As reported, E-selectin was upregulated in the vicinity of metastatic tumor cells in the lung, peaking approximately 6 hours after tumor cell inoculation and gradually decreasing thereafter.^21^ Accordingly, control and fucosylated OT-I cells (CD45.1+/-) were independently transferred into recipient mice with E0771-OVA lung metastasis at 6h after the intravenous injection of the tumor cells. Organs were harvested after 24 hrs for T cell quantification (Figure 3C). Among all the organs examined, only the lung was found to host a preferential infiltration of fucosylated OT-I T cells compared to control OT-I cells with twofold higher numbers of fucosylated cells observed (Figure 3D, Figure S4A). When adoptive transfer was conducted on day 7 after the lung metastasis had been established (Figure S4B), fucosylated OT-I cells were not only two to threefold higher than control OT-I cells in the lung (Figure S4C-D), but also slightly higher in other organs (Figure S4C), which further demonstrated that the increased tumor-specific homing induced by fucosylation is highly dependent on the E-selectin expression in tumors. Taken together, enforced fucosylation/display of sLe^x^ on the cell surface not only significantly enhanced T cell colonization in solid tumors (Figure 1) with high E-selectin expression, but also in the metastasis.

Next, we assessed *in vivo* anti-tumor efficacy of anti-EGFR CAR-T cells (E5-CAR-T cells) with or enforced E-selectin ligand expression and compared it with that of the unmodified T cells in mice with concurrent orthotropic breast cancer and lung metastasis established using E0771-E5 tumor cells. On day 8 post-tumor cell inoculation, fucosylated and control E5 CAR-T cells (1×10^6^) were adoptively transferred into separated groups of recipient mice. Following transfer, Fuc-E5 CAR-T completely cleared the lung metastasis by day3 (Figure 3F-H), whereas E5 CAR-T required 6 more days to achieve the same outcome (Figure 3F-H). Furthermore, Fuc-E5 CAR-T cells also demonstrated superior efficacy in suppressing orthotopic tumor growth. By day 26, 3 out of 7 mice in the Fuc-E5 CAR-T treated group became tumor-free, while the first complete response (CR) was observed on day 35 in the E5-CAR-T group (Figure 3H-I, K). Overall, the Fuc-E5 CAR-T treatment achieved a CR rate of 71% (5 out of 7 mice), significantly outperforming the E5-CAR-T group, which achieved a CR rate of 42% (3 out of 7 mice) (Figure 3J-K).

### Enforced expression of E-selectin ligands via cell-surface fucosylation enhanced CAR-T cell efficacy against lymphoma and leukemia

CAR-T cell-based immunotherapies have shown curative potential in B-cell malignancies.^23^ However, high production costs associated with extensive expansion and dose-dependent side effects significantly limit their widespread application. Inflammation is a common feature of lymphoma, a blood cancer that originates in the lymph nodes or the spleen, which triggers the upregulation of E-selectin.^24^ Likewise, E-selectin is constitutively expressed in the bone marrow niche^8^, where leukemia originates. For these reasons, we hypothesized that enforced expression of E-selectin ligands on CAR-T cells could enhance their therapeutic efficacy against blood malignancies, with the potential to reduce the infusion dose and minimize related adverse events. To this end, we first tested the efficacy of fucosylated anti-mouse CD19 CAR-T cells in an A20 B cell lymphoma model. To establish this model, Balb/c mice were treated with cyclophosphamide to induce lymphodepletion one day (Day -1) prior to tumor cell inoculation.^25^ After five days (Day 5), 8×10^5^ CAR-T cells were transferred (Figure 4A). Compared to control unmodified CAR-T cells, fucosylated CAR-T cells significantly delayed tumor growth (Figure 4B-C) and prolonged mice survival (Figure 4D).

**Figure 4.**
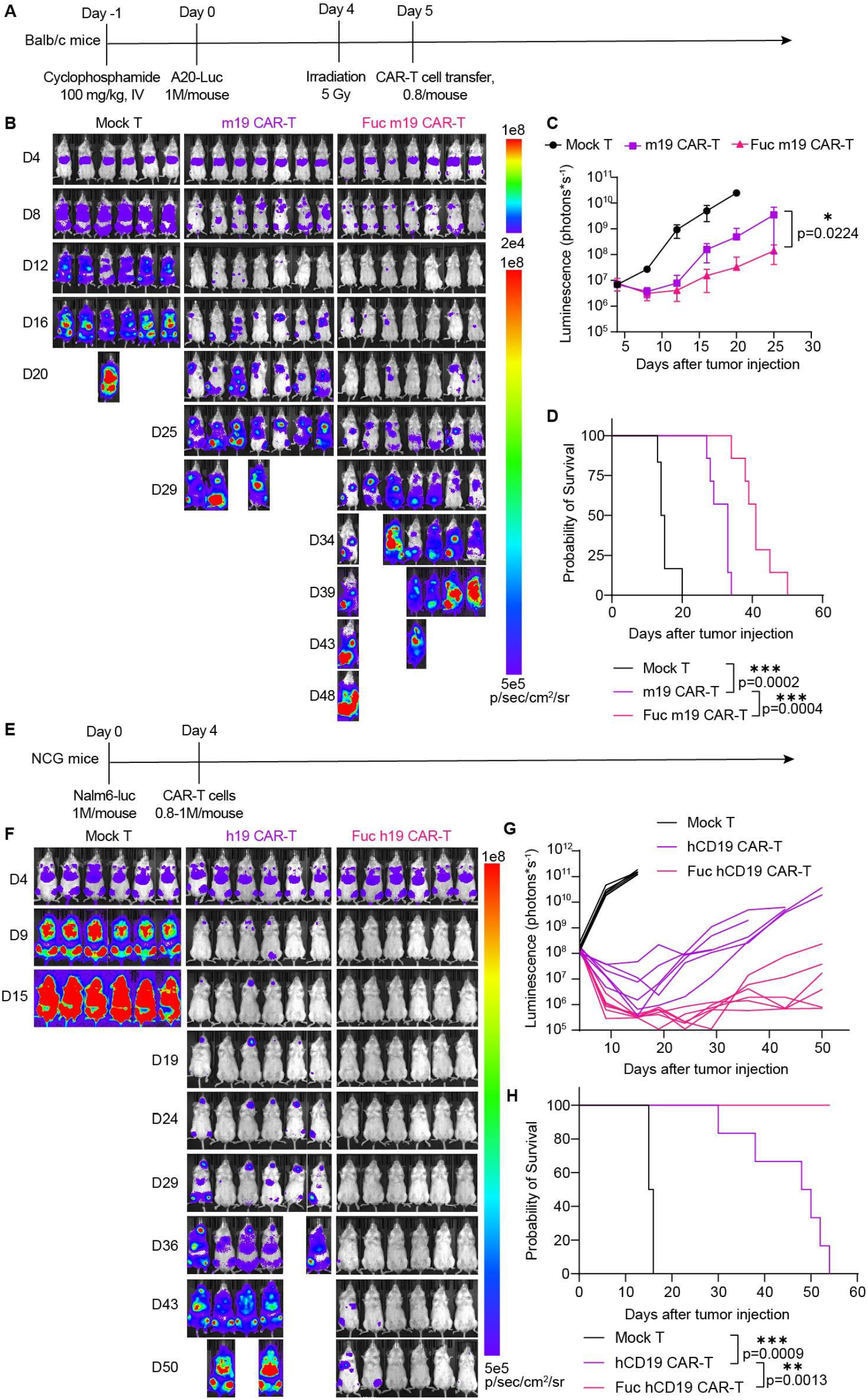
Cell-surface fucosylation on CAR-T cells improve the antitumor efficacy against mouse lymphoma and human leukemia. (A) Experimental setup. Balb/c mice were treated with cyclophosphamide (100 mg/kg, IV injection) and 1×10^6^ A20-luc cells were intravenously injected on the second day. Lymphodepletion was performed on day4. On day5, 0.8×10^6^ mock-transduced T cells or anti-CD19 CAR-T cells with or without fucosylation were transferred into tumor-bearing mice on (B-C) Tumor progression was monitored by IVIS and (D) mice survival was recorded. (E) 1×10^6^ Nalm6-luc cells were intravenously injected into NCG mice and 0.8-1×10^6^ anti-hCD19 human CAR-T cells with or without cell-surface fucosylation were transferred on day 4. (G) Tumor progression (front side of the mice) was monitored by IVIS and (H) mice survival was recorded. 6-7 repeats for each group (n=6-7), nsP > 0.05; *P < 0.05; **P < 0.01; ***P < 0.001; ****P < 0.0001; tumor size was analyzed by student T-test and survival curves were analyzed by log-rank test. Mean ±standard deviation (SD) values of biological replicates.

Building on the success with mouse CAR-T cells, we further evaluated the impact of enforced E-selectin expression on the therapeutic efficacy of human CAR-T cells in treating leukemia. Similarly to what we observed for fucosylated OT-I cells, fucosylated human anti-CD19 CAR-T cells exhibited higher levels of sLe^X^ (Figure S5A), and a slightly faster proliferation rate compared to unmodified CAR-T cells (Figure S5B). To understand the dynamics of enforced fucosylation, we used GF-biotin as the donor substrate to track the retention time of externally added fucose on CAR-T cell surface. As shown in Figure S5C-D, the level of biotinylated fucose decreased over time, but persisted for more than a week. Consistent with the findings in the mouse lymphoma model, significantly enhanced anti-leukemia activity was observed for fucosylated anti-human CD19 CAR-T cells in a xenograft human leukemia model. In this model, recipient mice were intravenously inoculated with Nalm6 expressing luciferase (Nalm6-luc) cells, a leukemia cell line derived from a patient with acute lymphoblastic leukemia. Adoptive transfer of conventional, unmodified anti-human CD19 CAR-T (0.8-1×1 0^6^) induced only transient tumor growth inhibition, and ultimately failed to control tumor progression (Figure 4F-H, Figure S5E-F). By contrast, all mice receiving fucosylated CAR-T cells achieved cancer-free status starting on day 9, consistent with findings in the murine lymphoma model. To day55, all mice treated with fucosylated CAR-T cells are still alive, with 3 out of 6 mice are tumor free while all mice treated with unmodified CAR-T cells were dead due to tumor burden.

### *In-situ* overexpression of human FUT6 impairs *in vivo* T cell survival

As E-selectin ligands added via FUT6-mediated cell-surface fucosylation decrease over time due to membrane protein turnover and cell division, we sought to overcome this limitation by overexpressing human fucosyltransferase 6 (Figure 5A) in the Golgi where the enzyme is naturally located. Overexpression of FUT6 in OT-I T cells had no negative impact on cell growth (Figure S6A). And the modified T cells exhibited significantly enhanced binding to E-selectin (Figure 5B) compared to control OT-I cells transfected with the empty viral vector (EV). However, a substantial portion of OT-I cells overexpressing FUT6 underwent apoptosis in the *in vitro* E-selectin binding assay (Figure S6B). Consistent with this finding, following adoptive transfer into either tumor-free or tumor-bearing mice (Figure 5C), very few OT-I cells overexpressing FUT6 could be detected in the blood or other organs 24h post-transfer (Figure 5D).

**Figure 5.**
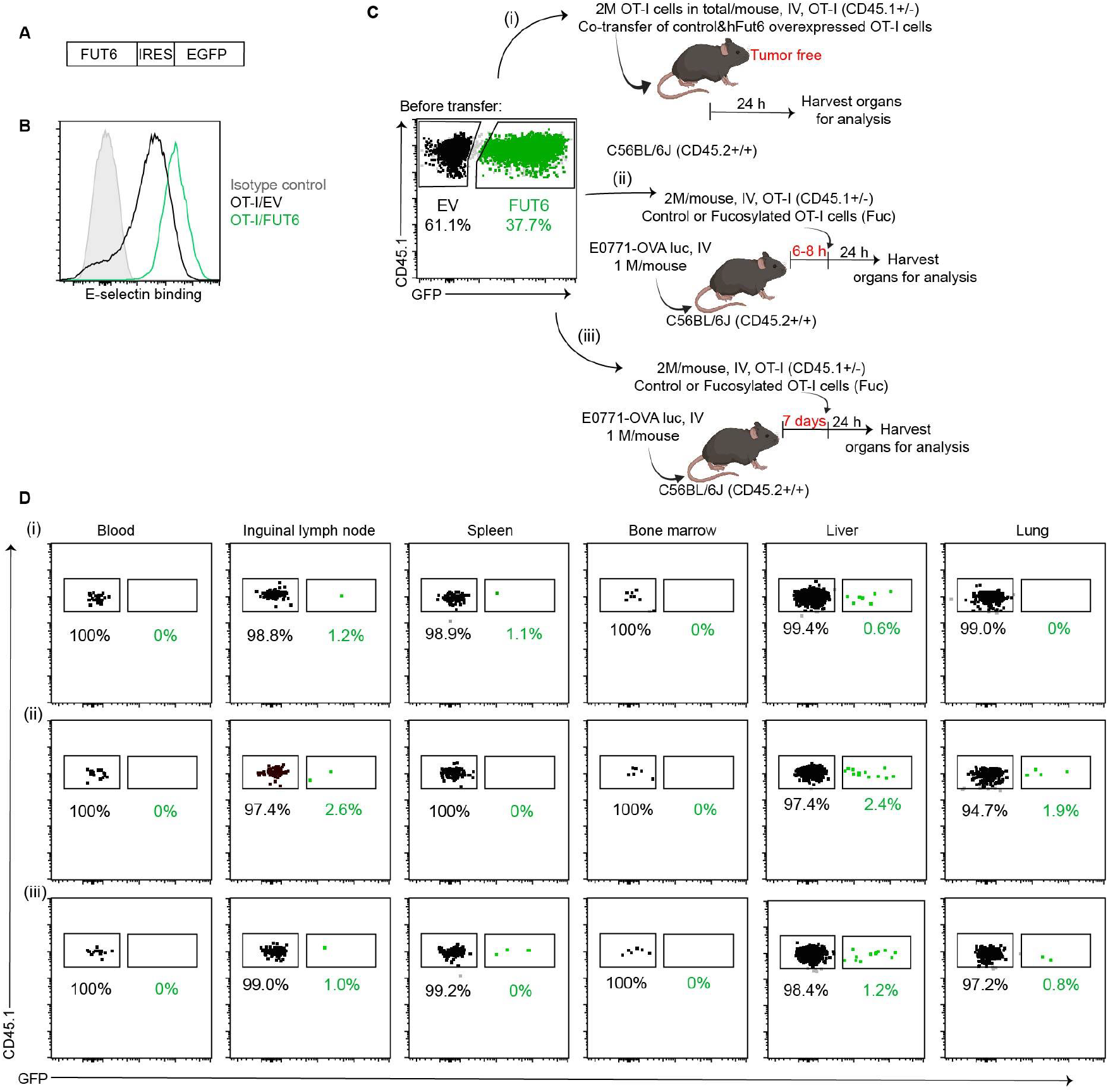
*In situ* overexpression of hFut6 impaired OT-I cells in vivo survival. (A) Construct of pMSCV-FUT6-IRES-EGFP, (B) *In vitro* E-selectin binding, and (C-D) *in vivo* homing of OT-I cells with or without overexpression of FUT6 in E0771-OVA tumor-free or tumor-bearing mice.

## Discussion

In this study, we performed the first comparison of enforced E-selectin ligand installation-achieved via FUT6-mediated direct cell-surface fucosylation versus FUT6 Golgi overexpression-on the *in vivo* homing and anti-tumor efficacy of genetically engineered T cells. We found that direct fucosylation significantly enhanced tumor-specific homing, which led to improved therapeutic efficacy of adoptive T-cell therapies. Notably, T cells modified by this approach exhibited increased expression of E-selectin ligands, which persisted for approximately 48 hours on murine T cells and up to 7 days on human T cells.

We also found for the first time that in tumor-free mice, T cells modified by direct fucosylation exhibited enhanced infiltration into various tissues without specificity, indicating an overall increase in migratory capabilities. By contrast, in mice with tumor burdens, these modified T cells preferentially migrated to E-selectin-expressing lesional tissues within 24 hours of adoptive transfer, and thereafter colonized the tumor parenchyma. Significantly, this high initial recruitment/parenchymal infiltration of T cells translated into improved anti-tumor efficacy across diverse cancer models, including bone marrow-originated tumors such as leukemia, lymphoid malignancies like lymphoma, solid tumors, and their lung and bone marrow metastases. Conversely, tumors with negligible E-selectin expression, such as B16 melanoma, showed no enhancement in T-cell homing or therapeutic benefit, highlighting the essential role of endothelial E-selectin expression in driving tissue specific T-cell colonization. To our surprise, efforts to achieve sustained sLe^X^ expression via FUT6 overexpression were counterproductive, in that resultant E-selectin ligand expression yielded reduced T-cell viability upon engagement with E-selectin. These findings, therefore, establish direct enzymatic cell-surface fucosylation as a simple and effective strategy to enhance tumor-specific targeting in adoptive T-cell therapies, particularly for cancers characterized by upregulated E-selectin expression.

In contrast to our findings, a previous study using human FUT7 to enforce sLe^X^ display on CAR-T cells indicated that while increased sLe^X^ surface expression enhanced E-selectin binding *in vitro*, but did not improve bone marrow homing or leukemia control *in vivo*^.17^ Notably, this study used a higher dose of CAR-T cells for adoptive transfer (3 million cells) compared to the 0.7-1 million T cells used in our experiments. The effect of cell dose on tissue saturation is a plausible explanation for this discrepancy^.14^ Intravenous infusion of high cell doses likely leads to saturation of tissue beds in a stochastic manner, thus, resulting in non-specific accumulation of administered cells independent of molecular effectors of migration. Conversely, at lower infusion doses, such as 1 million cells per mouse, the tissue parenchyma remains unsaturated, allowing molecular effectors, such as the newly added sLe^X^, to direct CAR-T cell colonization preferentially to tissues whose endothelial beds express E-selectin.

In particular, achieving the desired therapeutic effect with lower dosing offers several practical advantages. First, it has the potential to reduce the costs associated with T-cell expansion and mitigate T-cell exhaustion during expansion. It may also minimize the adverse effects associated with high-dose infusions.^4^ Taken together, these advantages may broaden the applicability and improve the safety profile of ACT, making it more accessible and versatile for the treatment of a wider range of diseases beyond cancer, such as inflammation and autoimmunity.

## Acknowledgement

This work was supported by the NIH (R35GM139643 to P.W., U01 CA225730 to. R. S.).

## Author contributions

Y. H. and P. W. conceived the project. Y. H., R.S. and P. W. designed experiments. J. Y. made all fucose derivatives. K. M. provided the human fucosyltransferase FT6. Y. H., K. Q., L. C., S. C., W. W., and M. C. performed the *in vivo* and *in vitro* experiments. Y. H., R.S. and P. W. analyzed and interpreted results. Y. H., R.S. and P. W. wrote the manuscript. All authors reviewed and edited the final version of the manuscript.

## Materials and Methods

**Table 1.**
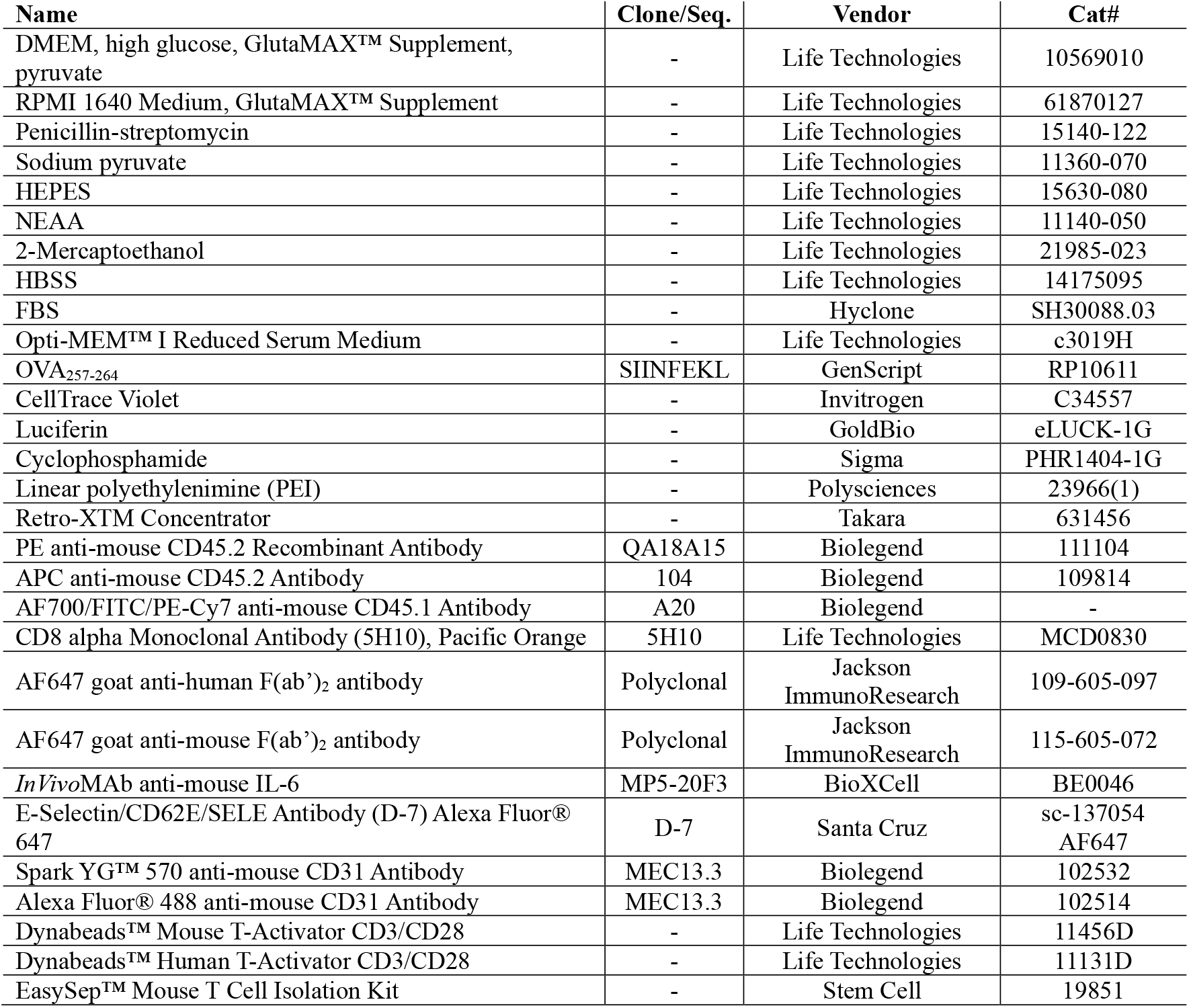
Reagents and antibodies.

### Mice

C57BL/6J (both WT and CD45.1+/+ congenic strains) mice were purchased from Jackson Laboratory. Balb/c mice were purchased from Charles River Laboratory. OT-I mice (strain B6.129S6-Rag2tm1FwaTg(TcraTcrb)1100Mjb, CD45.2+/+) were purchased from Taconic Biosciences. NOD-Prkdcem26Cd52Il2rgem26Cd22/NjuCrl (NCG) mice were purchased from Charles River. OT-I+/-CD45.1+/-mice were bred in-house using C57BL/6J (CD45.1+/+) and OT-I mice (CD45.2+/+). All the mouse study was started at the age of 8-10 weeks of the mice. All mice were bred and housed in specific pathogen free (SPF) rooms. All animal studies were approved by TSRI Animal Care and Use Committee.

### Cell lines

Cell lines were purchased from ATCC unless otherwise specified. E0771-OVA cells (provided by Dr. Zachary C.

Hartman at Duke University), MC38hCEA (provided by Dr. Alexey V. Stepanov at Scripps), Platinum-E cells (provided by Prof. John R. Teijaro lab at Scripps), and HEK293 Phoenix Ampho Packaging Cells (provided by Prof. Andrew Ward at Scripps) were cultured in complete DMEM medium (Life Technologies, Cat#10569-010) supplemented with 10% fetal bovine serum (FBS, Hyclone, Cat#SH30088.03) and 1% penicillin-streptomycin (Life Technologies, Cat#15140-122). Nalm6-Luc cells (provided by Prof. Crystal L Mackall’s lab at Stanford University) were cultured in RPMI1640 medium (Life Technologies, Cat#61870-127) supplemented with 10% FBS and 1% penicillin-streptomycin. OT-I cells and CAR-T cells were culture in RPMI1640 medium (Life Technologies, Cat#61870-127) supplemented with 10% FBS, 1% penicillin-streptomycin, 1 mM sodium pyruvate (100×stock: Life Technologies#11360-070), 10 mM HEPES (100×stock: Life Technologies#15630-080), 1×NEAA (100×stock: Life Technologies#11140-050), 55 uM 2-Mercaptoethanol (1000×stock: Life Technologies#21985-023), and 60 IU/mL rhIL-2.

### Ethics statement

All human blood samples were purchased from the Scripps General Clinical Research Center (GCRC) with informed consent of anonymous healthy donors via Scripps Research’s Normal Blood Donor Service (NBDS) under a Scripps Institutional Review Board (IRB) protocol. All human samples were analyzed with the approval of local ethical committees, in accordance with local regulations and U.S. Common Rule. Written informed consent was obtained from each participant.

### Cell-surface fucosylation

OT-I splenocytes were activated with OVA_257-264_ (1 nM, SIINFEKL, GenScript) for 2 days and expanded in T cell medium for 2-4 days. Then, OT-I cells were collected and resuspended in fresh T cell medium containing 0.5 mM GDP-Fucose and 0.05 mg/mL human fucosyltransferase 6 at a cell density of 10 M/mL. Fucosylation was performed in a 37 °C cell culture incubator for 30-60 min and cells were washed three times with T cell medium for further analysis. Fucosylation on CAR-T cells was performed following the same method as described above.

### *In vitro* E-selectin binding assay

Buffer 1: HBSS buffer containing 5%FBS, 2mM CaCl_2_

Buffer 2: HBSS buffer containing 5%FBS

0.5 million cells were stained with E-selectin-hFc (final concentration: 10 ug/ml) in 50 uL buffer 1 on ice for 20-30 min and washed with buffer 2 for three times. Then the cells were stained with APC anti-human Fc in buffer 2 on ice for 30 min and analyzed by flow cytometry after washed with buffer 2.

### *In vivo* distribution of OT-I cells or CAR-T cells

*E0771-OVA solid tumor model* 1×10^6^ E0771-OVA cells were implanted into the fourth mammary fat pad of female C57BL/6J mice (CD45.2+/+). Control OT-I cells (non-fucosylated, CD45.1+/-) were labeled by CellTrace Violet (CTV, Invitrogen, Cat.# C34557) and mixed with fucosylated OT-I cells (CD45.1+/-) at a ratio of 50%:50%. Then, the cells were co-transferred to the tumor bearing mice at tumor size around 100 mm^3^ (about day12-20 after tumor inoculation). 24h after the adoptive transfer, mice were euthanized and tissues including blood, lung, bone marrow, spleen, tumor draining lymph node, and tumor, were harvested and homogenized to single cell suspension. OT-I cells were identified by staining CD8, CD45.1, and CD45.2.

*E0771-OVA lung metastasis tumor model* 1×10^6^ E0771-OVA expressing luciferase cells were intravenously injected (tail vein) female C57BL/6J mice (CD45.2+/+). At 6h or on day 7 after the tumor injection, control and fucosylated OT-I cells (CD45.1+/-) were separately transferred by intravenous injection to the tumor bearing mice. At 24h after the adoptive transfer, tissues were collected for flow cytometry analysis as described above.

*E0771-OVA bone marrow metastasis tumor model* 1×10^6^ E0771-OVA expressing luciferase cells were injected to female C57BL/6J mice (CD45.2+/+) through caudal arteries. On day 7 after the tumor injection, control and fucosylated OT-I cells (CD45.1+/-) were separately transferred by intravenous injection to the tumor bearing mice. At 24h after the adoptive transfer, tissues were collected for flow cytometry analysis as described above.

*MC38hCEA tumor model* 1×10^6^ MC38hCEA tumor cells were subcutaneously implanted to the right flank of male or female C57BL/6J mice. Control anti-hCEA mouse CAR-T cells were labeled by CTV following the manufacturer’s instructions and mixed with fucosylated anti-hCEA mouse CAR-T cells at a ratio of 50%:50%. Then, the cells were co-transferred to the tumor bearing mice at tumor size around 100 mm^3^ (about day12-20 after tumor inoculation). At 24h after the adoptive transfer, tissues were collected for flow cytometry analysis. CAR-T cells were identified by GFP expression or staining with AF647 labelled anti-human F(ab’)_2_ antibody.

### *In vivo* extravasation analysis of CAR-T cells

*Flow cytometry analysis*: 1×10^6^ MC38hCEA tumor cells were subcutaneously implanted to the right flank of male or female C57BL/6J mice. 3×10^6^ anti-hCEA mouse CAR-T cells in total (CTV labelled control CAR-T cells mixed with fucosylated CAR-T cells at a ratio of 50%:50%) were co-transferred into the tumor bearing mice at tumor size around 100 mm^3^ (about day12-20 after tumor inoculation). 24h after the transfer, PE anti-mouse CD45.2 (3-5 ug/mouse) were intravenously injected into the mice 3 min before sacrifice. Then the tumors were harvested for flow cytometry analysis. *Immunofluorescence analysis*: 1×10^6^ MC38hCEA tumor cells were subcutaneously implanted to the right flank of male or female C57BL/6J mice. 1.5×10^6^/mouse fucosylated anti-hCEA mouse CAR-T cells were transferred into the tumor bearing mice at tumor size around 100 mm^3^. 24h after the transfer, the tumors were harvested for immunofluorescence analysis.

### *In vivo* anti-tumor study

*MC38hCEA model* 1×10^6^ MC38hCEA tumor cells were subcutaneously implanted to the right flank of male or female C57BL/6J mice. On day 8 after the tumor inoculation, the mice were irradiated with a dosage of 5 Gy for lymph depletion by irradiator (RS 2000 Small Animal Irradiator) 8-24h prior to T cell transfer, and 5×10^5^ Mock T cell, control or fucosylated CAR-T cells were intravenously injected to the mice. On day 14 and day 20 after CAR-T transfer, the mice were treated with anti-mIL-6 (150 ug/mouse, IV injected, BioXCell, Clone: MP5-20F3).

*E0771-E5 model* 1×10^6^ E0771-E5 tumor cells were implanted into the fourth mammary fat pad of female C57BL/6J mice and simultaneously 1×10^6^ E0771-E5 luciferase expressing cells were intravenously injected to the same mice. The mice were irradiated with a dosage of 5 Gy for lymph depletion by irradiator 8-24h prior to T cell transfer, and 1×10^6^ Mock T cells, control or fucosylated anti-EGFR mouse CAR-T cells were adoptively transferred to the tumor bearing mice on day 8 after tumor inoculation. The solid tumor size was recorded every two days, and the lung metastasis progression was measured by bioluminescent imaging using the IVIS imaging system. Briefly, 200 μl 15 mg/mL D-luciferin, potassium salt (GoldBio, Cat#eLUCK-100) was injected i.p. into each mouse and mice were imaged after 10 min using the IVIS imaging system (PerkinElmer). Values were analyzed using Living Image software. Mice were humanely euthanized when the mice have 15% of weight loss due to lung metastasis burden or tumor size is over 1000 mm^3^.

*A20 model* Balb/c mice were treated with cyclophosphamide (Sigma, 100 mg/kg, I.V.) and 1×10^6^ A20 expressing luciferase cells were I.V. injected at 24 after the treatment. The mice were irradiated with a dosage of 5 Gy for lymph depletion by irradiator 12-24h prior to T cell transfer, and 1×10^6^ Mock T cells, control or fucosylated anti-EGFR mouse CAR-T cells were adoptively transferred to the tumor bearing mice on day 5 after tumor inoculation. Tumor progression was measured following the same method as described above.

## Generation of murine constructs and retroviral transduction

DNA construct of fucosyltransferase or chimeric antigen receptor (CAR) was ordered from Twist Bioscience or generated in-house by PCR and directionally cloned into the pMSCV-IRES-GFP vector. The Plat-E RV packaging cells were plated into a 10-cm dish 24h prior to the transfection. The medium was changed to fresh medium (9.0 mL) when the cell density reached 50-70% of confluence. The cells were incubated in the new medium for 1-2h for transfection. For transfection, linear polyethylenimine (PEI, 25 kDa, PolySciences, 30 ug) suspended in Opti-MEM (500 uL, Life Technologies) was added into the plasmid (10 ug) suspended in Opti-MEM, incubated for 30 min at room temperature in the dark, and added dropwise into the dish of cells. 6-8 h after transfection, the medium was replaced with fresh medium (10.0 mL), and supernatants were collected at 36-40 h and 60-64 h after transfection. The retroviral vector in supernatants was concentrated using Retro-XTM Concentrator (Takara) following the manufacturer’s instructions and stored at -80 °C for further use.

For fucosyltransferase overexpression, OT-I splenocytes were activated with 10 nM OVA_257-264_ for 24h and transduced with pMSCV-FUT6-IRES-GFP retrovirus. The cells were expanded in T cell medium with 60 IU/mL rhIL-2 2-3 days for further study.

For CAR-T cell preparation, mouse CD3 T cells were isolated using a CD3 T cell isolation kit and activated with Dynabeads™ Mouse T-Activator CD3/CD28 for T Cell Expansion and Activation at 2:1 beads:cell ratio in T-cell medium. The cells were transduced with CAR-T retrovirus at 24-36h after the activation and maintained at 0.5-1×10^6^ million/mL for 2-3 days for further study.

## Human CD19 CAR-T cell preparation and *In vivo* evaluation of the anti-tumor activity for CAR-T cells against leukaemia

Cryopreserved human PBMCs were thawed and activated the same day with Dynabeads™ Human T-Activator CD3/CD28 for T Cell Expansion and Activation at 1:1 beads:cell ratio in T cell medium containing 150 IU/mL of rhIL-2. T cells were transduced with anti-CD19 lentiviral vector at 24-36h after the activation and maintained at 0.5×10^6^-1×10^6^ cells/mL in T cell medium with 150 IU/mL of rhIL-2. Transduction efficiency was determined by an anti-mouse IgG F(ab’)_2_ (AF647, Jackson ImmunoResearch, Cat#115-605-072) antibody.

Immunocompromised NOD-Prkdcem26Cd52Il2rgem26Cd22/NjuCrl (NCG) mice were purchased from Charles River. Six-to-eight-week-old female mice were intravenously inoculated with 1×10^6^ Nalm6-luc cells on day 0. 0.8-1×10^6^ Mock T cells or CAR-T cells were injected intravenously on day 4 after the tumor inoculation. Leukaemia progression was measured by bioluminescent imaging using the IVIS imaging system. Values were analyzed using Living Image software. Mice were humanely euthanized when mice demonstrated signs of morbidity and/or hind-limb paralysis.

## Statistics

All figures are representative of at least three experiments unless otherwise noted. All graphs report mean ±standard deviation (SD) values of biological replicates. The statistical significance of two-group comparisons was calculated using Student’s t-test. P<0.05 was considered significant and is designated with an asterisk in all figures.

## Reference

1 Skovgard, M. S. et al. Imaging CAR T-cell kinetics in solid tumors: Translational implications. Mol Ther-Oncolytics 22, 355–367, doi:10.1016/j.omto.2021.06.006 (2021).

2 Ying, Z. T. et al. Distribution of chimeric antigen receptor-modified T cells against CD19 in B-cell malignancies. BMC cancer 21, doi:ARTN 198 10.1186/s12885-021-07934-1 (2021).

3 Di, M. et al. Costs of care during chimeric antigen receptor T-cell therapy in relapsed or refractory B-cell lymphomas. Jnci Cancer Spect 8, pkae059, doi:10.1093/jncics/pkae059 (2024).

4 Rotte, A. et al. Dose-response correlation for CAR-T cells: a systematic review of clinical studies. Journal for immunotherapy of cancer 10, doi:ARTN e005678 10.1136/jitc-2022-005678 (2022).

5 Turtle, C. J. et al. CD19 CAR-T cells of defined CD4 : CD8 composition in adult B cell ALL patients. J Clin Invest 126, 2123–2138, doi:10.1172/JCI85309 (2016).

6 Nolz, J. C., Starbeck-Miller, G. R. & Harty, J. T. Naive, effector and memory CD8 T-cell trafficking: parallels and distinctions. Immunotherapy 3, 1223–1233, doi:10.2217/Imt.11.100 (2011).

7 Phillips, M. L. et al. Elam-1 Mediates Cell-Adhesion by Recognition of a Carbohydrate Ligand, Sialyl-Lex. Science 250, 1130–1132, doi:DOI 10.1126/science.1701274 (1990).

8 Schweitzer, K. M. et al. Constitutive expression of E-selectin and vascular cell adhesion molecule-1 on endothelial cells of hematopoietic tissues. American Journal of Pathology 148, 165–175 (1996).

9 Pober, J. S. et al. Two distinct monokines, interleukin 1 and tumor necrosis factor, each independently induce biosynthesis and transient expression of the same antigen on the surface of cultured human vascular endothelial cells. The Journal of Immunology 136, 1680–1687, doi:10.4049/jimmunol.136.5.1680 (1986).

10 Bevilacqua, M. P., Pober, J. S., Mendrick, D. L., Cotran, R. S. & Gimbrone, M. A. Identification of an Inducible Endothelial Leukocyte Adhesion Molecule. P Natl Acad Sci USA 84, 9238–9242, doi:DOI 10.1073/pnas.84.24.9238 (1987).

11 P Mann, A. & Tanaka, T. E-selectin: Its Role in Cancer and Potential as a Biomarker. Translational Medicine 01, doi:10.4172/2161-1025.s1-002 (2012).

12 Munro, J. M. et al. Expression of Sialyl-Lewis X, an E-Selectin Ligand, in Inflammation, Immune Processes, and Lymphoid-Tissues. American Journal of Pathology 141, 1397–1408 (1992).

13 Silva, M., Fung, R. K. F., Donnelly, C. B., Videira, P. A. & Sackstein, R. Cell-Specific Variation in E-Selectin Ligand Expression among Human Peripheral Blood Mononuclear Cells: Implications for Immunosurveillance and Pathobiology. Journal of immunology 198, 3576–3587, doi:10.4049/jimmunol.1601636 (2017).

14 Mondal, N., Silva, M., Castano, A. P., Maus, M. V. & Sackstein, R. Glycoengineering of chimeric antigen receptor (CAR) T-cells to enforce E-selectin binding. Journal of Biological Chemistry 294, 18465–18474, doi:10.1074/jbc.RA119.011134 (2019).

15 Sackstein, R. The First Step in Adoptive Cell Immunotherapeutics: Assuring Cell Delivery via Glycoengineering. Frontiers in immunology 9, doi:Artn 3084 10.3389/Fimmu.2018.03084 (2019).

16 Alatrash, G. et al. Fucosylation Enhances the Efficacy of Adoptively Transferred Antigen-Specific Cytotoxic T Lymphocytes. Clinical Cancer Research 25, 2610–2620, doi:10.1158/1078-0432.CCR-18-1527 (2019).

17 Sánchez-Martínez, D. et al. Enforced sialyl-Lewis-X (sLeX) display in E-selectin ligands by exofucosylation is dispensable for CD19-CAR T-cell activity and bone marrow homing. Clin Transl Med 11, doi:ARTN e280 10.1002/ctm2.280 (2021).

18 Yu, W. H. et al. Chemoenzymatic Measurement of LacNAc in Single-Cell Multiomics Reveals It as a Cell-Surface Indicator of Glycolytic Activity of CD8 T Cells. J Am Chem Soc 145, 12701–12716, doi:10.1021/jacs.3c02602 (2023).

19 Christofori, G. New signals from the invasive front. Nature 441, 444–450, doi:10.1038/nature04872 (2006).

20 Esposito, M. et al. Bone vascular niche E-selectin induces mesenchymal-epithelial transition and Wnt activation in cancer cells to promote bone metastasis. Nature cell biology 21, 627–+, doi:10.1038/s41556-019-0309-2 (2019).

21 Hauselmann, I. et al. Monocyte Induction of E-Selectin-Mediated Endothelial Activation Releases VE-Cadherin Junctions to Promote Tumor Cell Extravasation in the Metastasis Cascade. Cancer Res 76, 5302–5312, doi:10.1158/0008-5472.CAN-16-0784 (2016).

22 Kuchimaru, T. et al. A reliable murine model of bone metastasis by injecting cancer cells through caudal arteries. Nat Commun 9, doi:Artn 2981 10.1038/S41467-018-05366-3 (2018).

23 Cappell, K. M. & Kochenderfer, J. N. Long-term outcomes following CAR T cell therapy: what we know so far. Nat Rev Clin Oncol 20, 359–371, doi:10.1038/s41571-023-00754-1 (2023).

24 Totsuka, T. et al. Invasive human T lymphoma cells produce a novel factor that upregulates expression of adhesion molecules on endothelial cells.

25 Kueberuwa, G., Zheng, W. M., Kalaitsidou, M., Gilham, D. E. & Hawkins, R. E. A Syngeneic Mouse B-Cell Lymphoma Model for Pre-Clinical Evaluation of CD19 CAR T Cells. Jove-J Vis Exp, doi:ARTN e58492 10.3791/58492 (2018).

