## supplementary figures for "Enforced E-selectin ligand installation enhances homing and efficacy of adoptively transferred T cells"

Address: MB 208 10550 North Torrey Pines Road, La Jolla, California 92037,

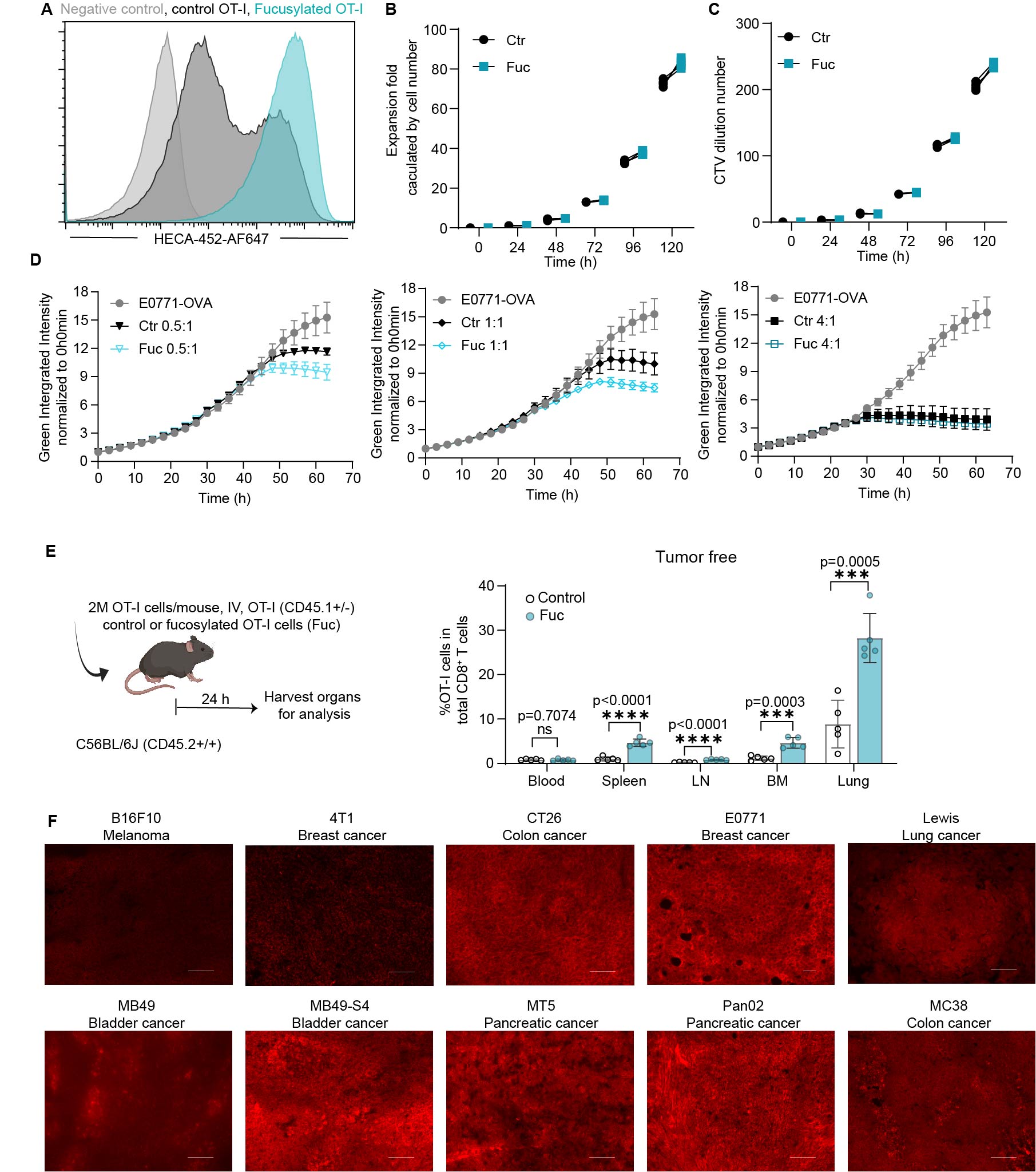

**Figure S1** (A) sLe^x^ on OT-I cell surface detected by a monoclonal antibody (HECA-452-AF647). OT-I cell expansion calculated by cell number (B) and CTV dilution number (C). *In vitro* anti-tumor activity of OT-I cells with (Fuc) or without (Ctr) fucosylation in different effector/tumor cell ratios as indicated in the figure (D). (E) OT-I cell ratios in total CD8^+^ T cells in different organs in tumor free mice. (F) E-selectin expression in tumors, scale bar: 100 µm. 3 repeats for in vitro and 5 repeats for in vivo studies in each group (n=3 or 5), nsP > 0.05; *P < 0.05; **P < 0.01; ***P < 0.001; ****P < 0.0001; analyzed by student T-test and mean ±standard deviation (SD) values of biological replicates.

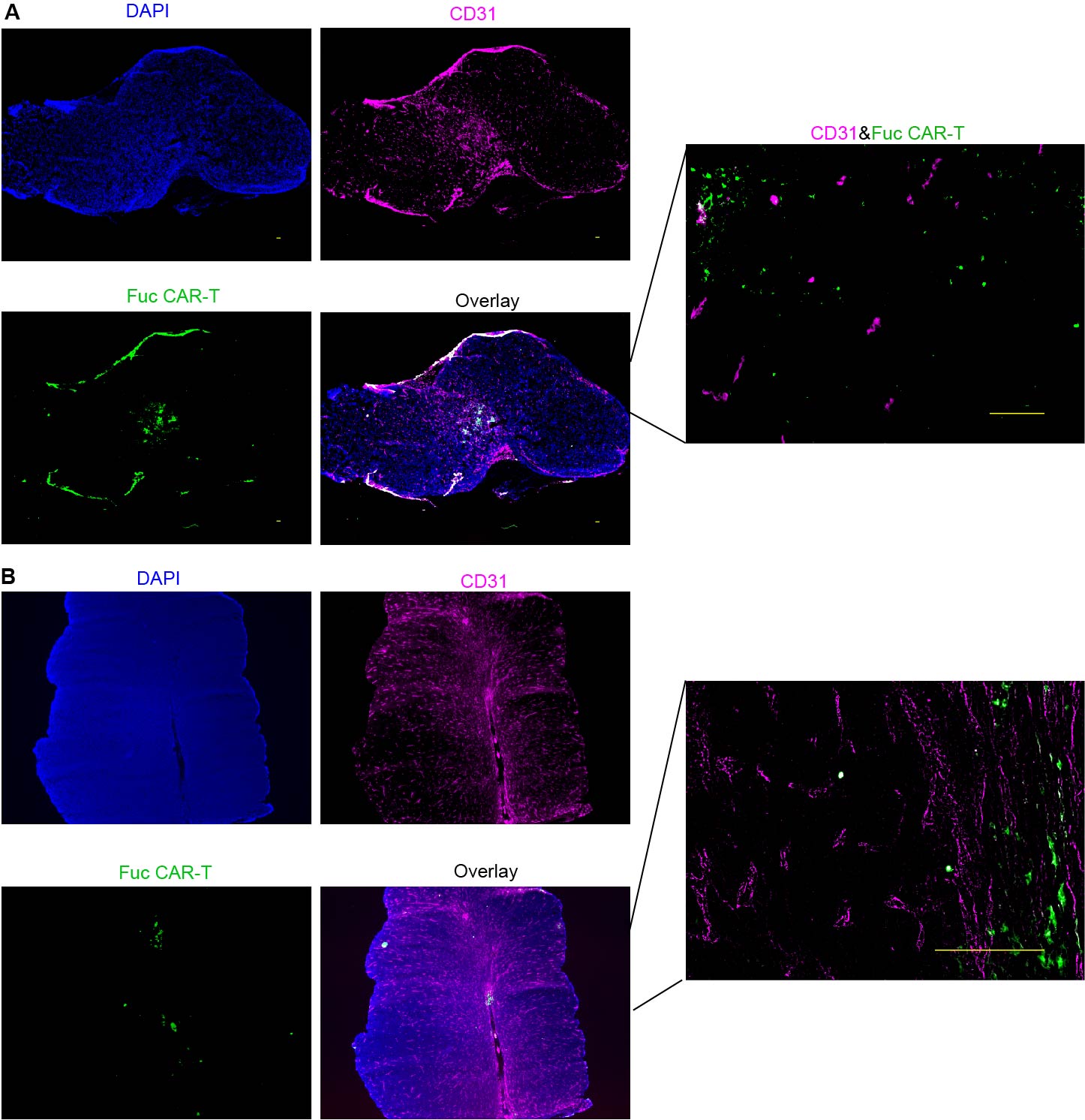

**Figure S2** Distribution of fucosylated CAR-T cells in the MC38hCEA tumor analyzed by immunofluorescence, tumor #2 (A) and tumor #3 (B), scale bar: 100 µm.

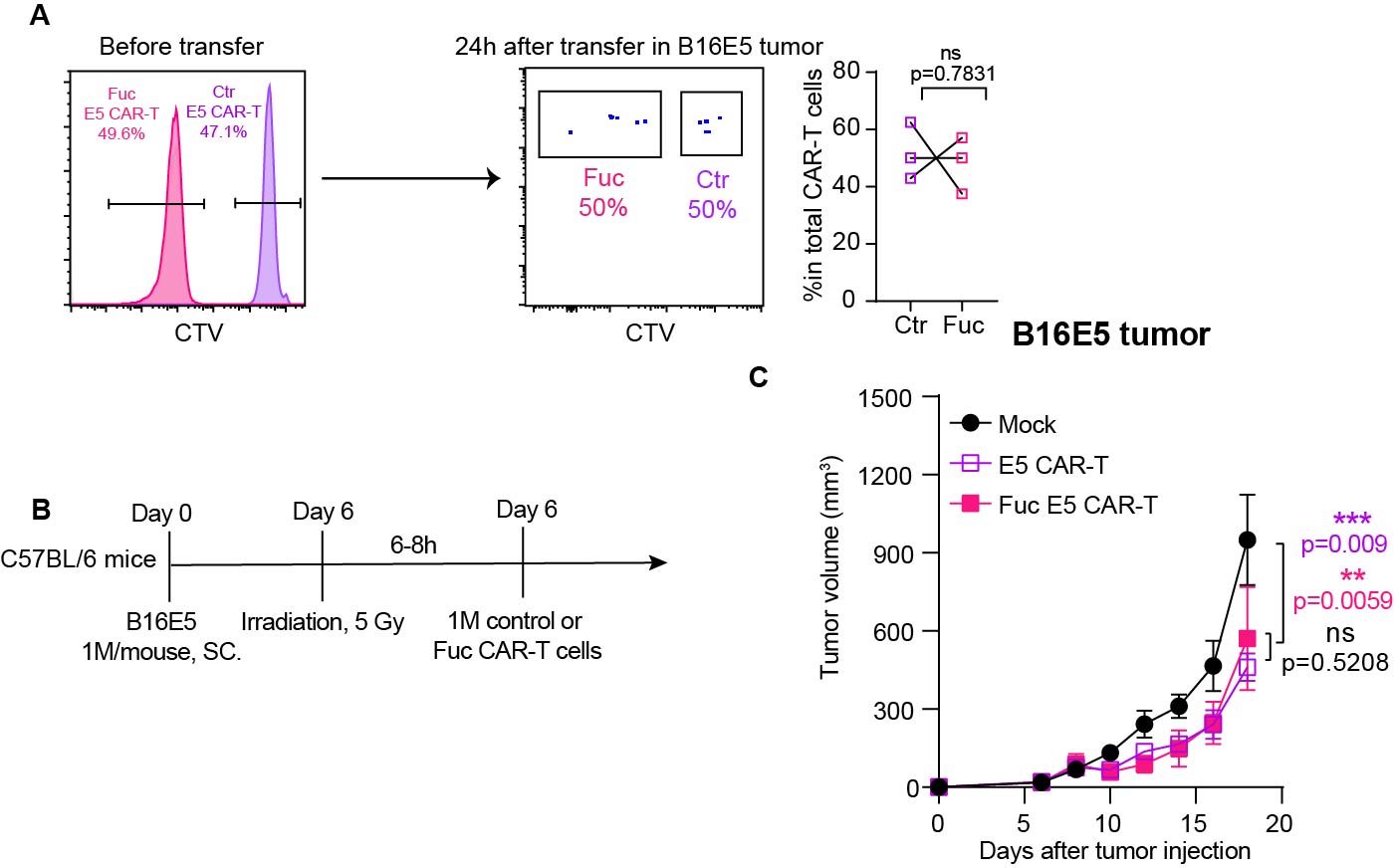

**Figure S3** **Cell-surface** **fucosylation on CAR-T cells doesn’t improve the tumor homing and control in B16F10 melanoma model.** (A) 1×10^6^ B16F10 expressing human EGFR (B16E5) cells were injected to subcutaneous right flank (Day 0) and 2×10^6^ CAR-T cells composed of unmodified E5 CAR-T cell (CTV positive) and fucosylated E5 CAR-T cells (CTV negative cells) in a 50%:50% cell ratio were transferred on day 8. Tumors were harvested at 24h after the CAR-T cell transfer for analysis. (B) 1×10^6^ B16F10 expressing human EGFR (B16E5) cells were injected to subcutaneous right flank (Day 0) and 1×10^6^ mock T cells or E5 CAR-T cells with or without fucosylation were intravenously injected to the tumor-bearing mice following lymphodepletion. Tumor growth was monitored every two days (C). 3-5 repeats in each group (n= 5), nsP > 0.05; *P < 0.05; **P < 0.01; ***P < 0.001; ****P < 0.0001; analyzed by student T-test and mean ±standard deviation (SD) values of biological replicates.

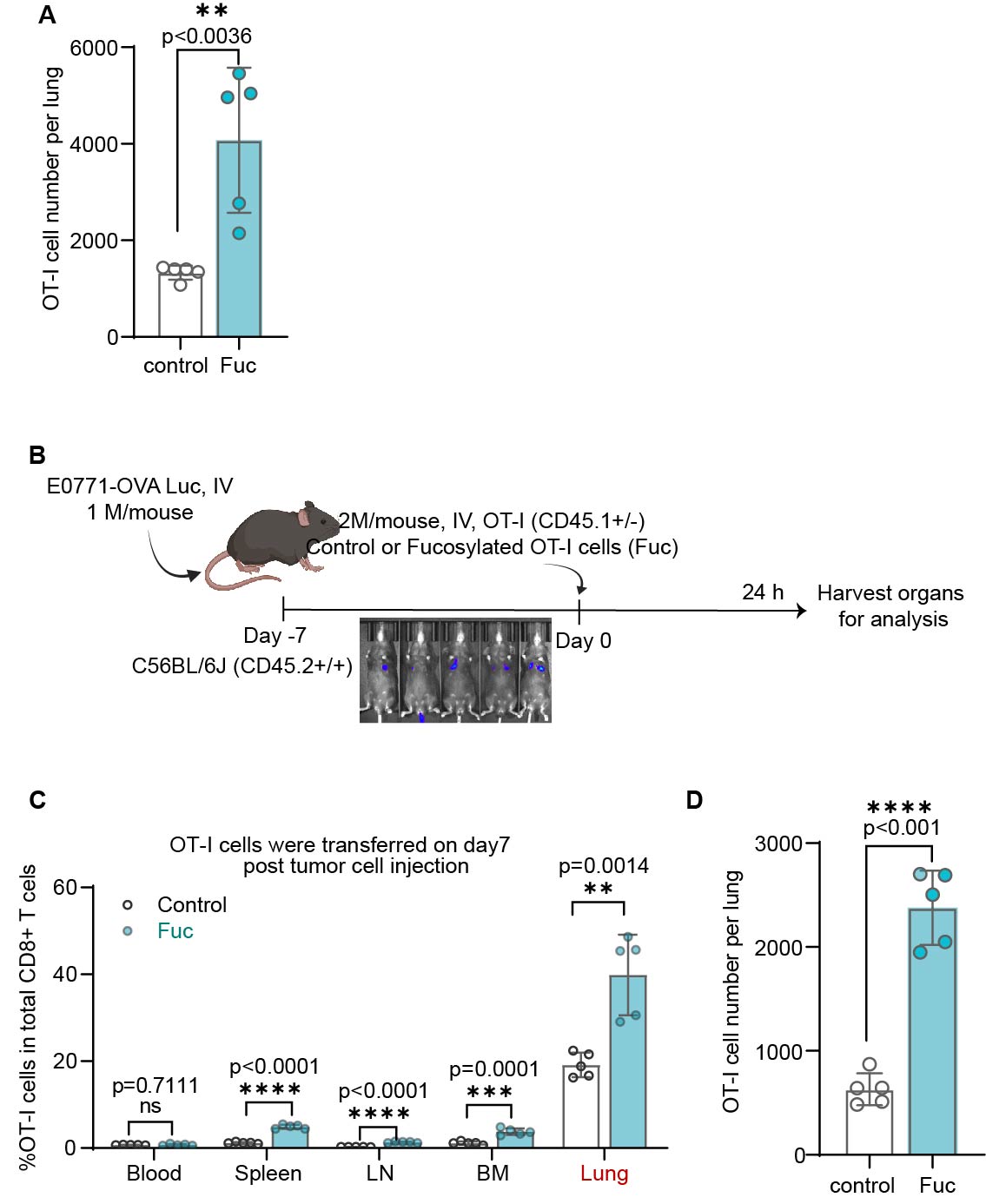

**Figure S4** **Cell-surface fucosylation increased CAR-T cells homing to lung metastasis.** (A) OT-I cell number in the lungs when OT-I cells were transferred at 6-8h after tumor cell injection. (B) 1×10^6^ E0771-OVA cells were intravaneously injected into the mice to develop lung metastasis and control or fucosylated OT-I cells were transferred on day 7 after tumor injection, (C) OT-I cell ratios in different organs were analyzed at 24h after the cell transfer and (D) OT-I numbers in the lungs. 3-5 repeats in each group (n= 5), nsP > 0.05; *P < 0.05; **P < 0.01; ***P < 0.001; ****P < 0.0001; analyzed by student T-test. Mean ±standard deviation (SD) values of biological replicates.

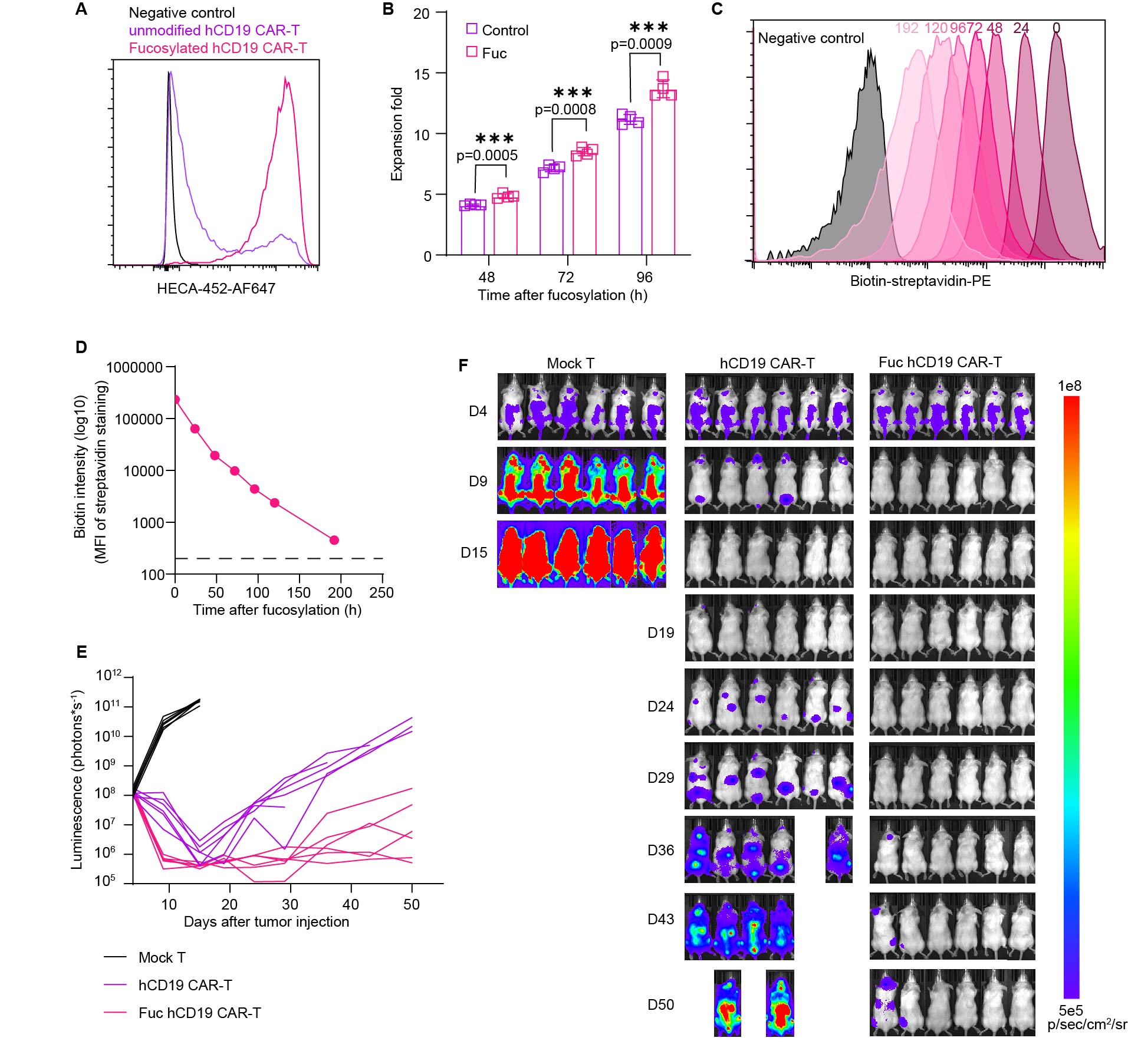

**Figure S5 *In vitro* characterization of cell-surface fucosylation on human anti-CD19 CAR-T cells and the *in vivo* anti-tumor efficacy.** (A) sLeX expression and (B) proliferation of human anti-CD19 CAR-T cells. (C-D) the decay of cell-surface fucosylation over time. (E-F) 1×10^6^ Nalm6-luc cells were intravenously injected into NCG mice and 0.8-1×10^6^ anti-hCD19 human CAR-T cells with or without fucosylation were transferred on day 4. Tumor progression (back side of the mice) (E) was monitored by. 4 repeats for in vitro assay and 6-7 repeats for in vivo experiment each group, nsP > 0.05; *P < 0.05; **P < 0.01; ***P < 0.001; ****P < 0.0001; the results were analyzed by student T-test. Mean ±standard deviation (SD) values of biological replicates.

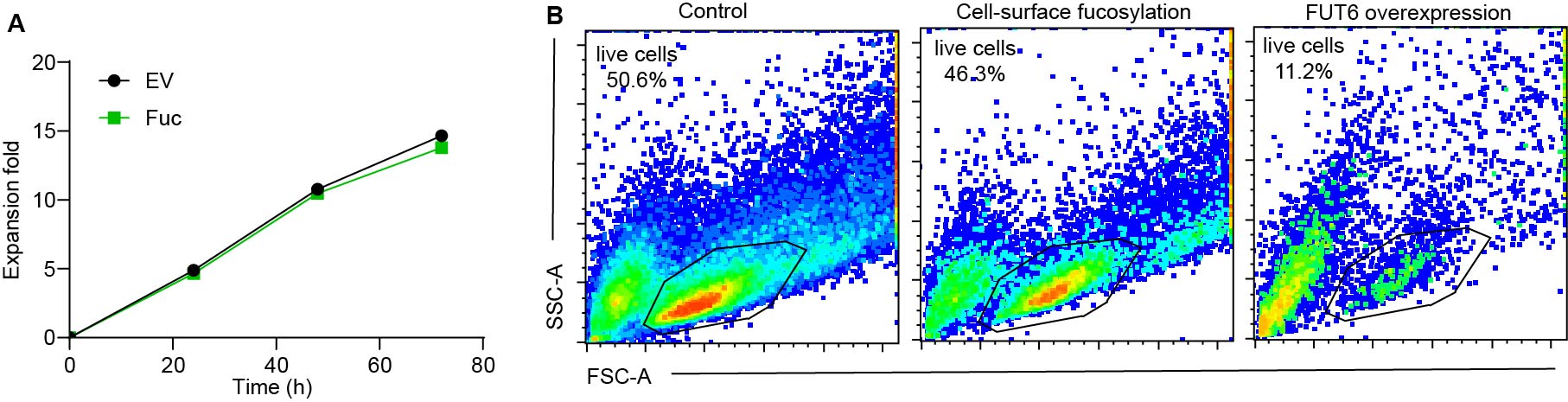

**Figure S6** (A) Proliferation of OT-I cells with or without FUT6 overexpression. (B) Live cell ratios of control OT-I cells, cell-surface fucosylated or FUT6 overexpressed OT-I cells after incubation with E-selectin. Mean ±standard deviation (SD) values of biological replicates.
